# Normal table of post-embryonic larval development for the California newt, *Taricha torosa*

**DOI:** 10.64898/2026.01.30.702904

**Authors:** Samantha M Westcott, Gary M Bucciarelli, Elizabeth A Heath-Heckman, Heather L Eisthen

## Abstract

We present here a normal table for post-embryonic development in the California newt (*Taricha torosa*), part of a genus of newts frequently studied for their toxicity and role within a predator-prey relationship. We generated the table by observing larvae collected as eggs in the wild and hatched and reared in the lab through metamorphosis. Building upon an established table consisting of 40 embryonic stages of development, our table consists of 13 stages based on discrete anatomical changes, primarily in limb development, and concludes at Stages 12-13 when the larvae undergo metamorphosis. We also describe more gradual phenotypic changes and their correlation to discrete stages in the developmental timeline. Finally, we illustrate the variability of the timing for reaching these stages in a controlled lab environment, demonstrating that time from hatching is not a reliable metric for standardizing results for diverse studies involving developing larvae. This staging table and accompanying observations will facilitate cross-study integration of research with larval *T. torosa*.

## Introduction

Many variables influence an animal’s development, including genetics, hormonal changes, nutrition, and environmental factors. Reliable developmental benchmarks enable a more complete understanding of a developing organism by facilitating the integration of information across labs and disciplines. Amphibians undergo extreme developmental changes post-hatching; these animals are also indicators for environmental health, and much research focuses on learning how development and survival may be affected by environments altered due to pollution [1], habitat disruptions [2], and climate change [3]. By documenting typical developmental patterns in a controlled environment, we can better understand potential deviations from these patterns.

Historically, much amphibian research has focused on embryonic development, starting from fertilized eggs. General embryonic development has been documented for frogs with a free-living larval stage in the Gosner staging table [4] which is based on development of the Gulf Coast toad, *Bufo valliceps*. The general model for embryonic development in salamanders is documented in Harrison’s staging table, which is based on spotted salamanders, *Ambystoma maculatum* [5]. Few staging tables describe the post-embryonic development of salamanders, including tables for *Ambystoma* spp. [6][7] and *Triturus* spp. [8] which provide descriptions of changes in pigmentation, presence of balancers, and order of digit development.

Twitty and Bodenstein [9] documented the embryonic development of California newts, *Taricha torosa*, in a table consisting of 40 distinct stages. Researchers seeking to describe biological processes in post-hatching larval *Taricha* typically either refer to the last few stages of Harrison’s staging table, which refers to salamanders from a different family, or simply state the time from hatching in an effort to standardize their assessment of larval development [10]. Experiments with wild-caught larvae in particular would benefit from a clear system to identify the developmental stage of larvae and facilitate comparisons with lab-based experiments. Here, we describe the post-embryonic development of *T. torosa* based on external characteristics and propose a uniform staging system to help standardize future research.

## Materials and Methods

### Research animals

We collected and housed clutches of *Taricha torosa* eggs at Harrison Stage 22 from a pond in Santa Rosa, California on 28 February 2024 under California Department of Fish and Wildlife Scientific Collection Permit #012430 GMB. Eggs and larvae were reared in an animal facility at Michigan State University. All experiments were approved by the MSU Institutional Animal Care and Use Committee (protocol approval nos. 201800106 and 202400136).

Eggs and larva were maintained at 19° ± 1.5°C on a 12:12 day:night cycle in glass embryo dishes filled with RO water fortified with salts and buffers (1.5 x 10^-4^% Replenish, 6.5 x 10^-5^% Stability, 3.8 x 10^-5^% Alkaline Buffer in RO water; Seachem Laboratories, Inc, Madison, GA, USA). We maintained aeration and water flow through the use of air stones and performed partial water changes daily.

The five larvae used in this study all hatched on 18 March 2024 in a dish housing three egg clutches. Upon hatching, each larva was moved to a separate dish. Larvae were fed daily with live brine shrimp (*Artemia sp*.) until large enough to be fed live blackworms (*Lumbriculus sp*.).

### Imaging equipment

We photographed each larva once a day for two months, at which time their external physical development slowed. We then gradually reduced our rate of imaging to once a week until metamorphosis or, in one case, mortality. For imaging, unanesthetized larvae were transferred to a Petri dish filled with buffered water. Larvae were gently maneuvered under a microscope using a transfer pipet. Images of the dorsal side of each animal were collected through metamorphosis; images of the ventral side were collected until the animal no longer tolerated being inverted.

Initial images were visualized through a Leica M125C stereo microscope collected using the Leica Application Suite X v5.2.0 and until the larvae grew too large to capture the main body at the lowest magnification. Larvae were then examined under a Zeiss Stemi 2000-C stereo microscope and images were collected with a mounted Nikon EOS Rebel T3 camera.

### Image editing

We modified raw images using Adobe Photoshop 2025 to optimize visual comparisons of the developing larvae and remove distracting background. Specifically, we conservatively deleted the backgrounds of the raw images but, to avoid deleting any portion of bodily structures, did not erase portions of background around fine features such as gill filaments, balancers, and digits. We also adjusted the brightness and contrast of images to create an even appearance across panels in assembled figures, but we did not adjust color. All such changes were applied uniformly across entire images and no edits were made to single portions of images. Thus, for example, artifacts such as *Artemia* eggs or fibers in the water that overlapped with any part of an animal’s body were not deleted from images and we did not modify glare or artifacts caused by photographing animals in water.

## Results

We recorded larval development from the initial day of hatching through the time the animals completed metamorphosis (July 2024) or mortality. In our staging system, described below, stages 1-12 are primarily marked by the appearance and development of limbs and digits, while Stage 13 depends largely on the reduction in gills. Larvae begin metamorphosing during Stage 12 with the completion of the process correlating with the end of Stage 13. Additionally, we describe other important physical changes that occur throughout these thirteen stages of development — more gradual changes that cannot readily be categorized as discrete stages. Finally, we document the variability in timing with which these discrete and gradual changes occurred. These descriptions provide a thorough account of the external morphological changes taking place during larval development in *T. torosa*.

### Discrete anatomical stages

We identified thirteen distinct developmental stages (Fig 1). For simplicity, our staging table begins at larval Stage 1 and is based on discrete changes, primarily in limb and digit formation. Due to variability in hatching rates, some larvae were already showing signs of developing digits on the forelimb upon hatching; thus, some individuals hatch at Stage 2 of our staging system.

**Figure 1.**
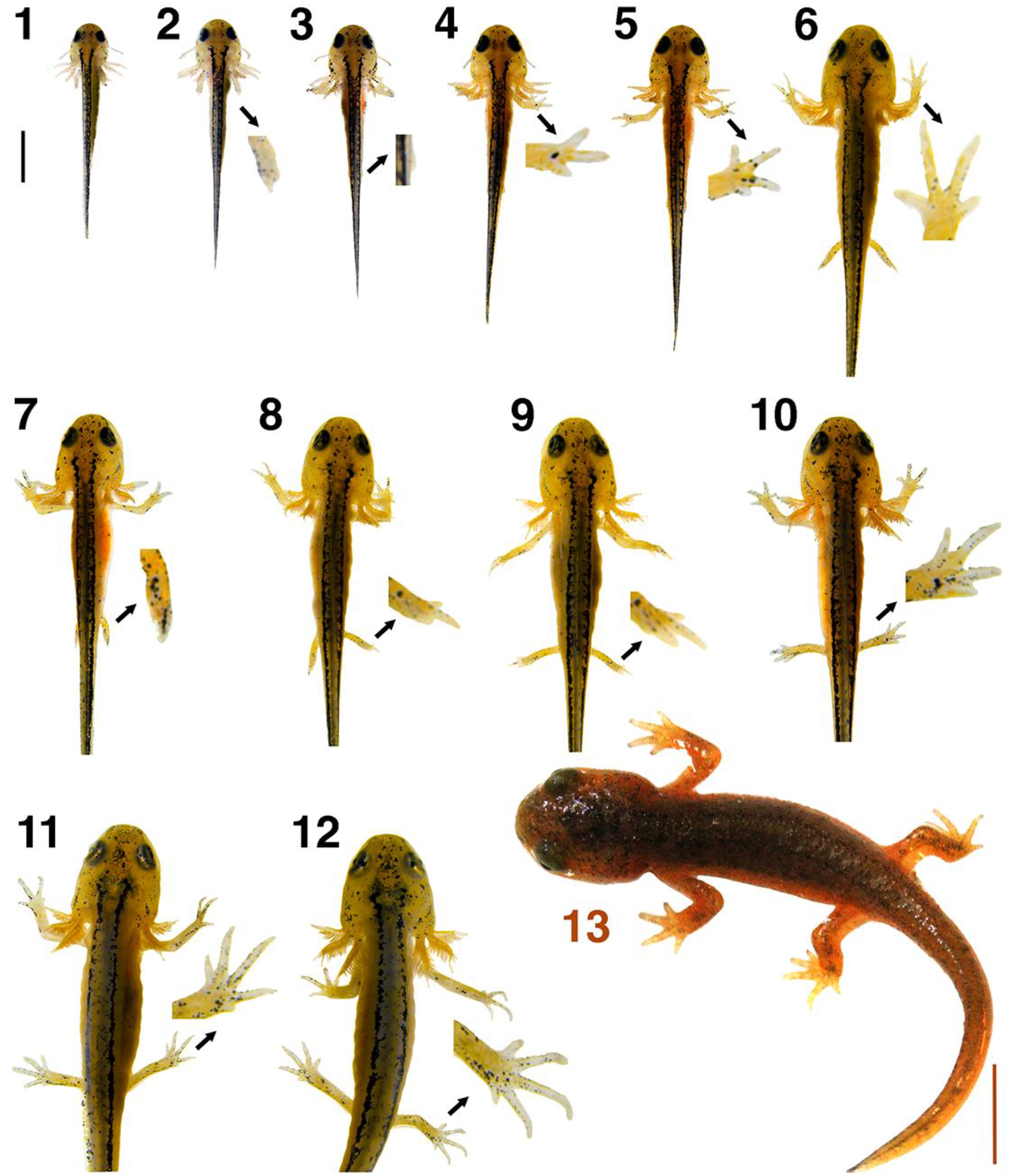
Post-hatching phenotypic changes for staging *T. torosa*. Each larva is labeled by stage number. To make fine details clear, stage 2-12 larvae are accompanied by an expanded image of a key feature used to identify their stage; these insets are enlarged 16.7x from the associated photo for Stages 2-10 and 8.3x for Stages 11-12. Black scale bar in the upper left corner (= 2 mm) applies to the images of Stage 1-12 larvae and the orange scale bar in the bottom right corner (= 5 mm) applies to the image of the Stage 13 larva.

### Stage 1: Forelimb buds present and elongating

*T. torosa* larvae hatch with forelimb buds and balancers present. The forelimbs begin to develop within the egg as small, rounded buds that then elongate dorsally, with the distal end of the buds flattening as they lengthen. The larvae also possess a visible yolk sac occupying much of the pleuroperitoneal cavity. Larvae did not eat for the first few days post-hatching.

### Stage 2: Initial digits emerging on forelimb

Distal ends of the forelimbs show signs of the first digits emerging as slight bifurcated projections.

### Stage 3: Hindlimb buds emerging

Hindlimb buds begin to form on the ventrolateral side of the larvae, slightly anterior to the cloaca. These limb buds are barely visible in a dorsal view and can more readily be seen from a lateral or ventral view. Meanwhile, the forelimbs lengthen, begin to bend, and show clear signs of cubital joint development.

### Stage 4: Three clear digits on forelimb

The digit primordia on the distal end of each forelimb have resolved into three digits. The two preaxial digits are long and roughly similar in length and a smaller postaxial digit primordium is visible.

### Stage 5: Balancers reabsorbed and fourth forelimb digit emerging

The balancers undergo apoptosis in early development of hatchlings, a process that is apparent at Stage 4 and completed by Stage 5. Simultaneously, the fourth and final digit primordium on each forelimb becomes visible, appearing in the postaxial position.

### Stage 6: Fourth forelimb digit clear

Four distinct digits are now present on each forelimb. Since the development of the third digit, the relative lengths of the digits have remained roughly the same. The hindlimbs are elongated and can now project laterally away from the body.

### Stage 7: First digits emerging on hindlimb

The first digits of the hindlimbs arise on the distal ends, with one clear central projection between subtle developing primordia on either side.

### Stage 8: Three digits on hindlimb

The initial digit primordia have resolved into three distinct digits on each hindlimb, with the middle digit remaining the longest.

### Stage 9: Fourth hindlimb digit emerging

A fourth digit primordium begins to form in the postaxial position on each hindlimb.

### Stage 10: Fourth hindlimb digit clear

The fourth hindlimb digits are now distinct.

### Stage 11: Fifth hindlimb digit emerging

The fifth and final hindlimb digit primordium begins to form as the last postaxial digit on each hindlimb. The central digit is now the longest, extending past the adjacent preaxial digit. A loose webbing of skin has arisen along the posterior ventral side of the hindlimbs.

### Stage 12: Fifth hindlimb digit clear

Stage 12 lasts much longer than any other stage, as larvae enter metamorphosis and begin to resemble juvenile newts. Because of the lack of subsequent discrete changes, we use skin texture changes to further subdivide Stage 12 into early, middle, and late substages, as discussed below. As illustrated in Figure 1, at the beginning of Stage 12, the fifth and final hindlimb digits are now distinct. The larvae still possess gills dense with secondary filaments showing no reduction in length as well as smooth skin and dorsal pigmentation patterns similar to those of hatchling larvae. Larvae undergo a prolonged period of growth in early Stage 12. Signs of metamorphosis first become noticeable mid-Stage 12: the skin is no longer smooth and the gills begin to recede. During late Stage 12, the skin develops clear bumps and the gills are nearly resorbed, with few remaining secondary filaments. More detailed descriptions of the gradual changes occurring in middle and late Stage 12 larvae are described below, particularly in the section on skin roughness.

### Stage 13: Gills have mostly receded and larvae are nearing the end of metamorphosis

The larvae are near the end of metamorphosis. External gill structures are almost completely gone, with only small remnants of the gill rami present near the base of the skull and no remaining secondary gill filaments. The skin is rough and larvae possess the dark brown and orange coloration characteristic of an adult newt. The tail fins have been completely resorbed.

### Gradual anatomical changes

Some features of the developing larvae undergo more nuanced changes that are difficult to categorize as distinct stages. These include changes in gill structure, pigmentation, and skin texture. We describe details of these developmental patterns here.

#### Skin becomes rough

The skin visibly changes in texture throughout Stage 12, and the progression of changes in skin texture can be used to denote how far into Stage 12 a larva has progressed. Fig 2A depicts the uniformly smooth skin texture that persists in larvae from Stage 1 through early Stage 12. After a prolonged period of larval growth in early Stage 12, the skin texture starts visibly changing, characterizing the substage that we refer to as mid-Stage 12 (Fig 2B). The surface of the skin now appears more complex, with slight bumps arising and branching iridophores visible along the dorsum. In late Stage 12 larvae, larger and more distinct bumps develop across the dorsal surface of the animal, giving the skin a rough texture (Fig 2C). At the apex of each bump is a visible pore and spot of black pigment makes the pores stand out against the rest of the skin coloration. By Stage 13, the iridophores are no longer visible along the dorsal surface of the skin (Fig 2D).

**Figure 2.**
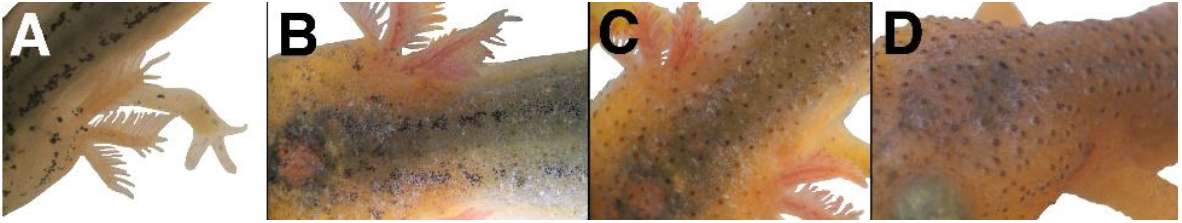
Gradual change in skin roughness. Close up images of the dorsum near the head of a single larva at different points in metamorphosis. Early in Stage 12 (**A**), the skin is smooth. By mid-Stage 12 (**B**), the skin becomes textured and iridophores become visible along the dorsal surface. In the late Stage 12 (**C**), raised bumps are present with black pigment at the peaks. The bumps persist beyond metamorphosis, but the iridophores are no longer visible by Stage 13 (**D**).

#### Gill filament density increases; gills then recede

As illustrated in Fig 3A, Stage 1 larvae possess external gills consisting of three paired rami. The rami are well established, with thick but sparse secondary filaments present. As the larvae grow, their gills become more elaborate. The rami expand in size and the secondary filaments increase in density. By Stage 6, the gill filaments populate most of the ventral surface area of the gill rami. The vasculature in the gills increases visibly, giving the filaments a red hue. Filament density peaks around Stage 9 (Fig 3B).

**Figure 3.**
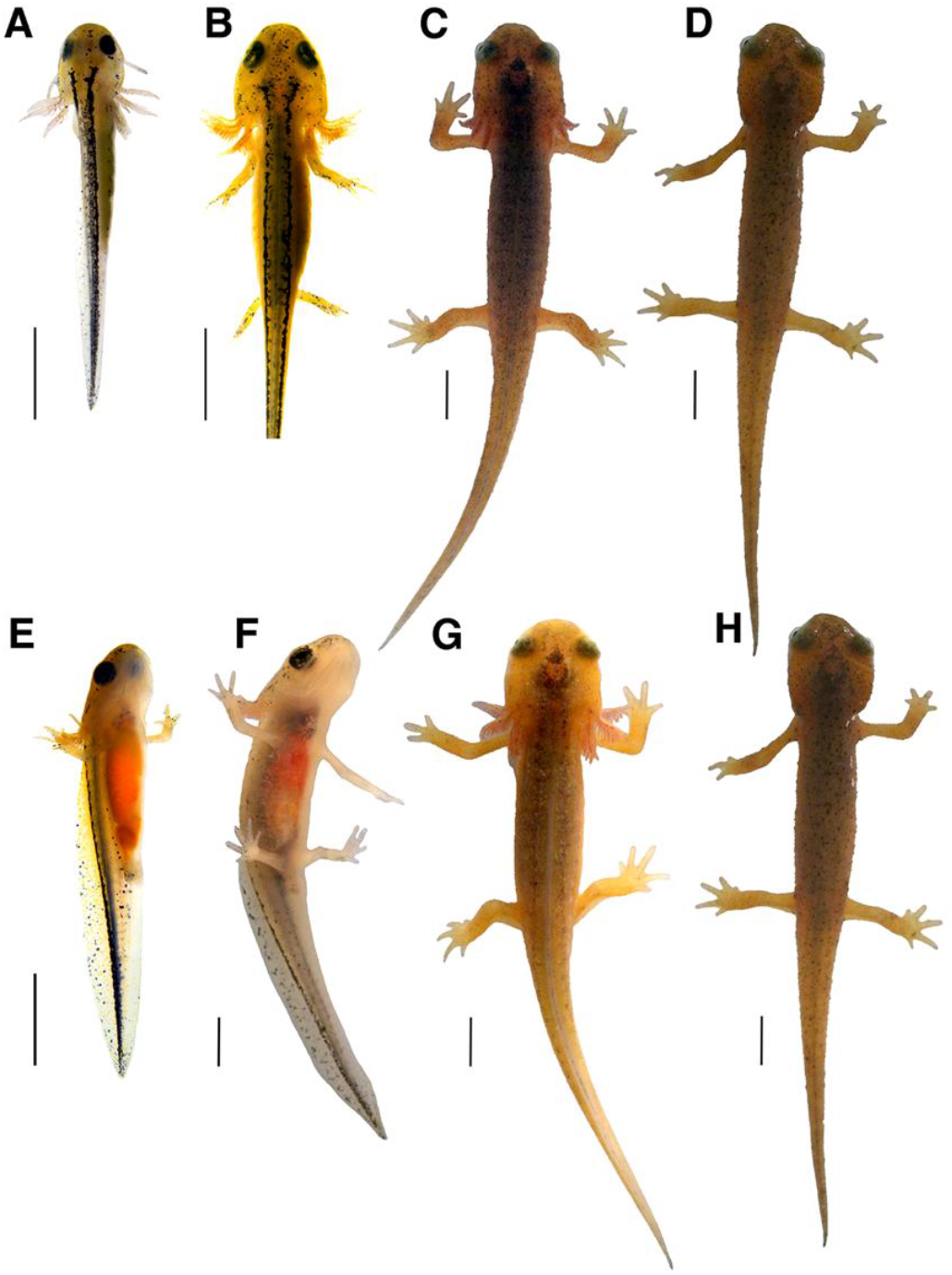
Gradual changes in gill (top) and fin (bottom) structures. The top row depicts the process of gill expansion and then retraction. At Stage 1 (**A**), the larva has long gill rami with few secondary filaments. These filaments increase in density and become fluffier, as seen in Stage 9 (**B**). Once the animal begins to metamorphose in Stage 12 (**C**), the gills begin to retract and secondary filaments are lost. The final portion of the receding gills can be seen at Stage 13 (**D**) as small nodules near the gular folds of the larva. The bottom row depicts the gradual loss of tail fins across larval development in lateral (**E** and **F**) and dorsal (**G** and **H**) views. At Stage 6 (**E**) the dorsal and ventral fins are at their peak size in proportion to the larva. The dorsal fin extends all the way from the tail tip to the just posterior of the head, while the ventral fin extends from the tail tip to the cloaca. The ventral fin decreases in height rapidly during forelimb development and is nearly absent by early Stage 12 (**F**), while the dorsal fin is slightly reduced in height. By mid-Stage 12 (**G**), the fin has receded posteriorly toward the tail and is greatly reduced in height before completely receding at Stage 13. All images are photographs of the same larva and all scale bars = 3 mm.

Secondary gill filaments reach their maximum density and length by Stage 12. The gills begin to recede during late Stage 12: the filaments decrease in number, and the rami shorten and recede towards the gular fold (Fig 3C). Once this process begins, the gills shrink dramatically over the course of a few days. At Stage 13 the rami project posteriorly and have receded to the point that only small nodules with no visible filaments remain (Fig 3D). The rami recede completely shortly after the final filaments.

#### Tail fins develop, then recede

Stage 1 hatchlings possess large, paddle-like tail fins that extend both ventrally and dorsally (Fig 3E). Starting at the tail tip, the dorsal fin projection extends anteriorly in a low arch and runs along approximately two thirds of the body length. The ventral fin extends from the distal end of the tail to the cloaca. Both fins are largely translucent but xanthophores are visible, particularly along the dorsal fin. Melanocytes are also present in low abundance and increase in density along the fin’s length as the larvae continue to develop. Overall, the pigmentation of the dorsal fin becomes noticeably denser around Stage 4. The ventral fin visibly decreases in height beginning at Stage 4, becoming flatter in shape but persisting throughout most larval development stages before receding during early Stage 12. Early in Stage 12 the anteriormost point of the dorsal fin begins to incrementally recede toward the posterior end of the larvae while also gradually decreasing in height (Fig 3F). Late Stage 12 larvae possess only small remnants of the dorsal fin along the tail (Fig 3G), and the fin fully recedes in Stage 13 (Fig 3H).

#### Dorsal stripes change in clarity

Around Harrison Stage 36 [11], embryos begin to develop two dorsal stripes of melanophores that run in parallel along either side of the vertebral column. Fig 4A illustrates an example of the dorsal stripes present at hatching. The pigment cells become more organized over time, and the lines become clearer and better-defined over much of larval development. Pigment also appears as large spots occupying the space medial to the two stripes. Both the lateral stripes and medial clusters are transient and defined stripes are no longer visible by the end of metamorphosis. The lateral stripes consist of superficial melanophores near the epidermis [12], while the melanophores forming the medial clusters appear to reside deeper in the dermis. The dissolution of the lateral stripes is accompanied by an increase in clarity of the medial clusters. As the larvae grow, the paired lateral stripes appear clearer due to the increasing fusion of the dermal pigment forming the medial clusters (Fig 4B). Concurrently, the lateral stripes become discontinuous, with gaps appearing between melanophores along the length of the stripes. By Stage 10, pigment at the anterior-most region of the lateral stripes dissipates (Fig 4C). The pigment cells continue to shrink in size until the stripes are completely absent. Fig 4D illustrates the final remnants of the dorsal stripes, in a mid-Stage 12 larva. Small melanophores appear sparsely along the dorsum, just visible against the maturing pigmentation typical of an adult.

**Figure 4.**
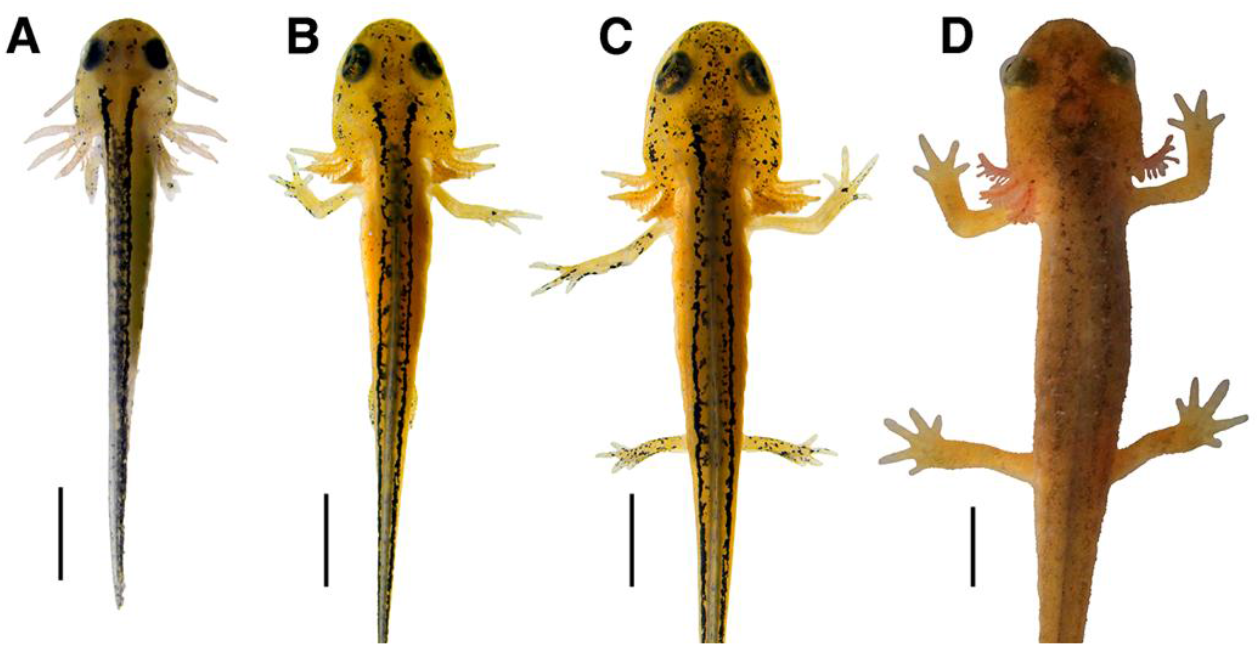
Gradual changes in dorsal stripes. Images of the progression of dorsal stripe definition over time. Early-stage larvae possess dorsal stripes consisting of tightly packed melanophores on either side of their dorsal fin, as seen in Stage 2 (**A**). The stripes become somewhat obscured by clusters of pigment in the medial space between the lateral stripes. The lateral stripes then grow thinner and small gaps appear as illustrated in a Stage 8 larva (**B**). As the animal grows, the medial pigment clusters become more diffuse before the lateral stripes dissipate at the anterior end in Stage 10 (**C**). The last remnants of the lateral stripes are visible at mid-Stage 12 and the medial clusters are obscured by the maturing dorsal pigmentation (**D**). All images are photographs of the same larva and all scale bars = 3 mm.

#### Ventral side becomes more opaque

As shown in Fig 5A, the ventral side of Stage 1 larvae is translucent. The contents of the pleuroperitoneal cavity are visible, which at this stage is primarily filled by a yellow-green yolk sac. The heart is also visible anterior to the yolk sac. The ventral sides of their large eyes are visible through the membrane separating the oral and optic cavities as well as the thin, unpigmented layers of skin and muscle composing their lower jaws. The yolk sac rapidly decreases in size over Stages 1-4. Fig 5B (Stage 10) illustrates the progression of the ventral tissue development as the skin grows more opaque, starting laterally and progressing towards the midline. Also visible at this stage are the intestines and liver, no longer hidden by the presence of the now-resorbed yolk sac. Simultaneously, the pericardium appears thicker and heavily pigmented, obscuring the visibility of the heart. Fig 5C illustrates an early Stage 12 larva in which the ventral skin is nearly opaque aside from a small patch along the midline. The entirety of the ventral surface becomes opaque around mid-Stage 12, after which the skin becomes pigmented, gaining the characteristic orange color of adult *Taricha* as the animal progresses through metamorphosis.

**Figure 5.**
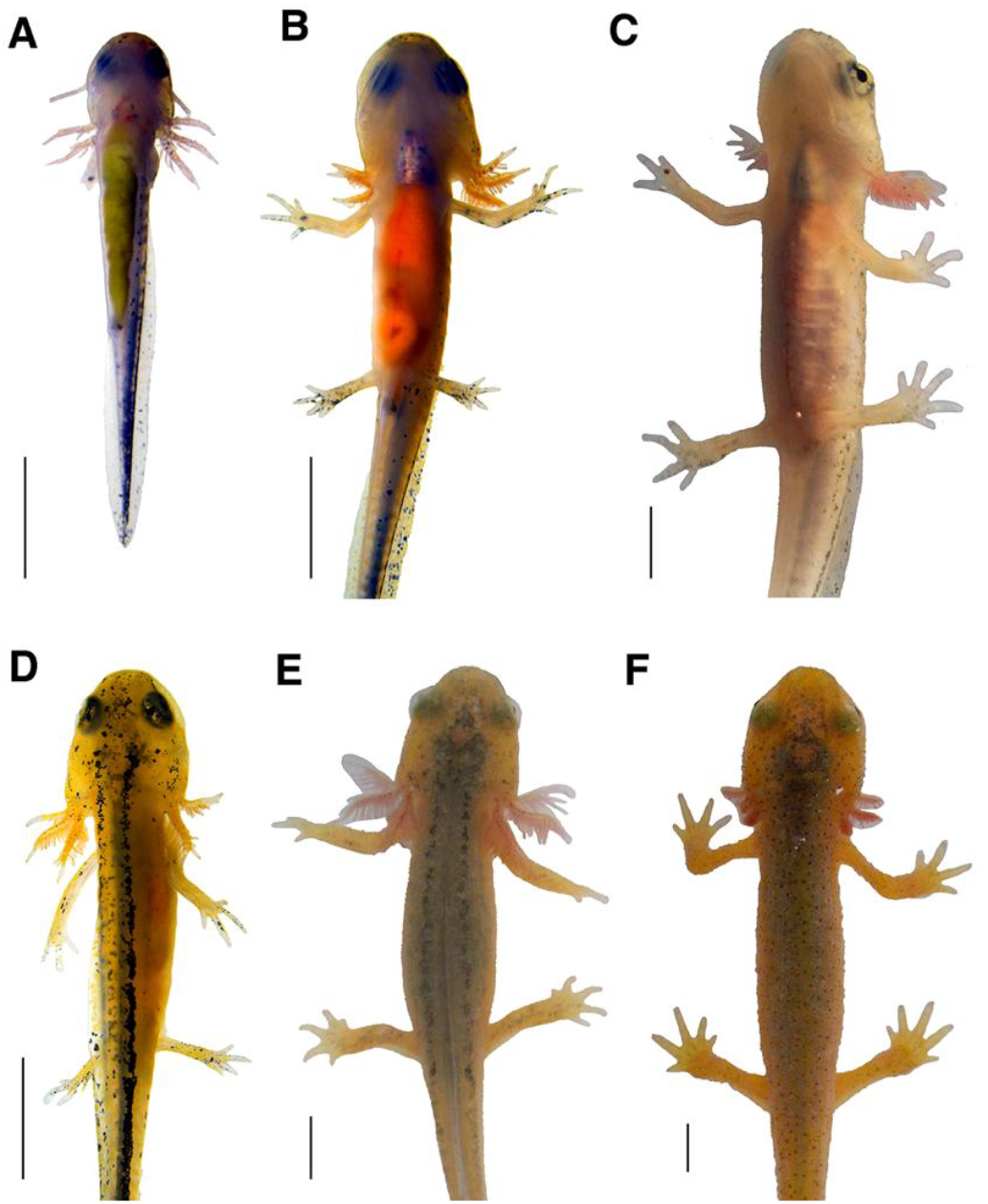
Gradual changes in ventral opacity and dorsal pigmentation. The top row depicts increasing ventral opacity in a single larva. At Stage 2 (**A**), several internal structures are clearly visible, including the yellow-green yolk that occupies most of the pleuroperitoneal cavity, the eyes, and the heart. By Stage 10 (**B**) the lateral skin is thickening and the heart is obscured by a thickening and pigmented pericardium. By early Stage 12 (**C**), the organs are just barely detectable through the skin. The bottom row shows the darkening in dorsal pigmentation in a different larva. At Stage 10 (**D**) the coloration of the dorsum is still largely unchanged since hatching. The color darkens gradually into an ashy brown color by mid-Stage 12 (**E**), darkening into a deep brown by late Stage 12 (**F**). All images are photographs of the same larva and all scale bars = 3 mm.

#### Dorsum becomes darker

Aside from stripes of melanophores, the dorsal skin appears largely unpigmented during much of larval development. The skin remains largely unpigmented until Stage 11 (Fig 5D). Changes in the coloration of the dorsum overlaps greatly with the dissolution of the dorsal stripes depicted in Fig 4. The skin color progresses from a light, ashy brown color arising at Stage 11 to a deep brown characteristic of mature *Taricha* at late Stage 12 (Fig 5F). Throughout larval development, the pineal organ is visible through the thin tissue layers forming the dorsal surface of the head. This forebrain structure is gradually enveloped by melanophores in the meninges as depicted in Figs 5E and F (mid-to-late Stage 12). The region overlying the pineal organ remains sparsely pigmented through the end of metamorphosis into the early juvenile stage, at which point the dorsum has fully darkened in color.

#### Eye pigmentation changes

Fig 6 illustrates development of larval eye pigmentation. Early post-hatching larvae possess large, uniformly black eyes that occupy a large proportion of the animal’s head (e.g. Fig 6A). Figs 6B and 6C illustrate the progressive accumulation of golden-brown pigment in the iris and dissipation of a black pigmented ring that surrounds the eye, a process first noticeable starting at Stage 6. Fig 6D illustrates the dark horizontal segmentation of the iris by early Stage 12, typical of a mature *T. torosa* eye.

**Figure 6.**
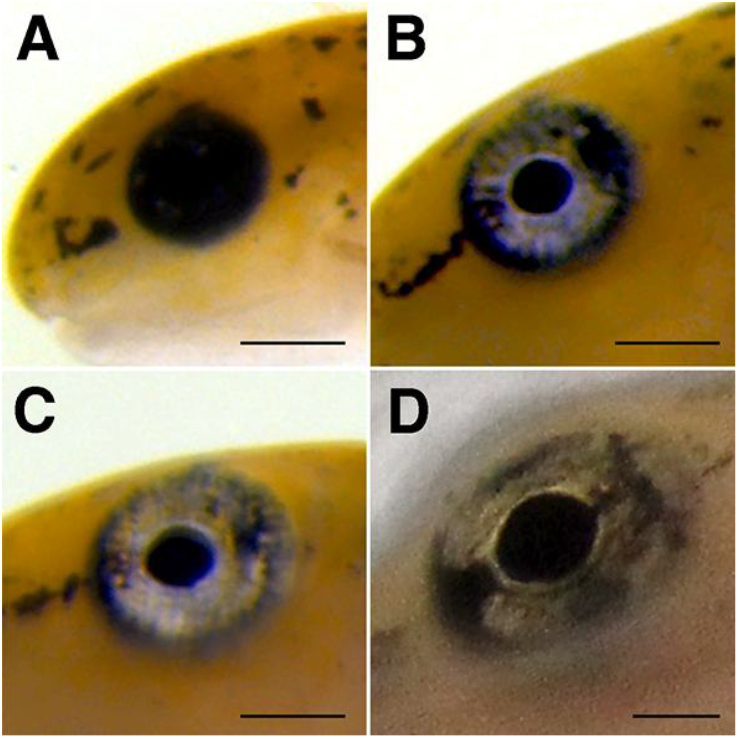
Gradual changes in eye pigmentation. Photographs illustrating changes in eye pigmentation of a single larva across time. (**A**) The eyes are primarily black with few discernable details in early larval stages, as seen in Stage 4. (**B**) By Stage 9, much of the black pigment has receded from the eyes, leaving a black ring of pigment surrounding the eyes and making the irises readily visible. By Stage 11 (**C**) most of the surrounding black pigment ring has dissipated and by early Stage 12 (**D**) has fully receded. At this stage, a clear horizontal bar bifurcates the golden irises. Scale bars = 500 µm.

### Variation in developmental rates

Developmental rates are not uniform under controlled laboratory conditions. Even with larvae originally collected as eggs from the same pond on the same day, hatched on the same day and reared under the same conditions, the time to reach each development stage was variable (Table 1). As noted earlier, larvae do not strictly hatch at the same developmental stage. Of the five larvae we observed, one hatched at our Stage 1, the equivalent of Harrison’s Stage 41, while four hatched at our Stage 2, equivalent to Harrison’s Stage 42. Differences in rates of development also varied among stages. This pattern of variable development rate persisted throughout the study.

**Table 1.** Days after hatching at which each larva reached each developmental stage.

| stage | morphology | Days after hatching |  |  |  |  | Average |
| --- | --- | --- | --- | --- | --- | --- | --- |
|  |  | Larva 1 | Larva 2 | Larva 3 | Larva 4 | Larva 5 |  |
| 1 | forelimb buds present and elongating | 0 | 0 | 0 | 0 | 0 | 0 |
| 2 | initial digits emerging on forelimb | 0 | 0 | 0 | 0 | 2 | 0.4 |
| 3 | hindlimb buds emerging | 3 | 3 | 6 | 7 | 8 | 5.4 |
| 4 | three clear digits on forelimb | 6 | 6 | 7 | 9 | 11 | 7.8 |
| 5 | balancers reabsorbed; fourth forelimb digit emerging | 17 | 17 | 16 | 17 | 20 | 17.4 |
| 6 | fourth forelimb digit clear | 18 | 18 | 18 | 18 | 21 | 18.6 |
| 7 | first digits emerging on hindlimb | 20 | 20 | 21 | 20 | 23 | 20.8 |
| 8 | three digits on hindlimb | 24 | 24 | 23 | 23 | 25 | 23.8 |
| 9 | fourth hindlimb digit emerging | 27 | 27 | 28 | 29 | 31 | 28.4 |
| 10 | fourth hindlimb digit clear | 33 | 33 | 34 | 34 | 36 | 34 |
| 11 | fifth hindlimb digit emerging | 38 | 38 | 39 | 38 | 42 | 39 |
| 12 | fifth hindlimb digit clear | 51 | 51 | 51 | 54 | 60 | 53.4 |
| 13 | gills mostly receded | 107 | 107 | 109 | 109 | * | 108 |

\* This individual died 115 days post-hatching, before reaching stage 13.

Figure 7 illustrates this variability by providing a comparison of two larvae (larvae 4 and 5 of Table 1) at hatching on March 18, 2024, as well as 28, 105, and 109 days after hatching. Fig 7A illustrates the slight differences between the larvae on their day of hatching: both larvae possess limb buds, but larva 4 (on the right) has more elongated buds with digit primordia present. Fig 7B depicts the same larvae 28 days after hatching; both larvae have reached Stage 8 but larva 4 is nearing Stage 9 and possesses fuller gills and has much longer hindlimbs. As demonstrated in Fig 7C, the differences are more dramatic 105 days after hatching. Larva 5 is at early Stage 12 while larva 4 is mid-Stage 12. Larva 4 is more developed, with the tail mostly absorbed, the gills shortened, the dorsal stripes much lighter, and overall pigment starting to resemble a mature newt. Meanwhile, the less developed larva 5 still looks very much like a juvenile. Finally, Fig 7D shows the more developed larva, larva 4, in the climax of metamorphosis, 109 days after hatching. Larva 4 has entered Stage 13, with only the smallest remnants of gills visible near the gular fold, rough, bumpy skin, a fully receded tail fin, and no dorsal stripes. Meanwhile larva 5 has finally entered mid-Stage 12 with slightly rougher skin, and no other notable phenotypic changes have occurred.

**Figure 7.**
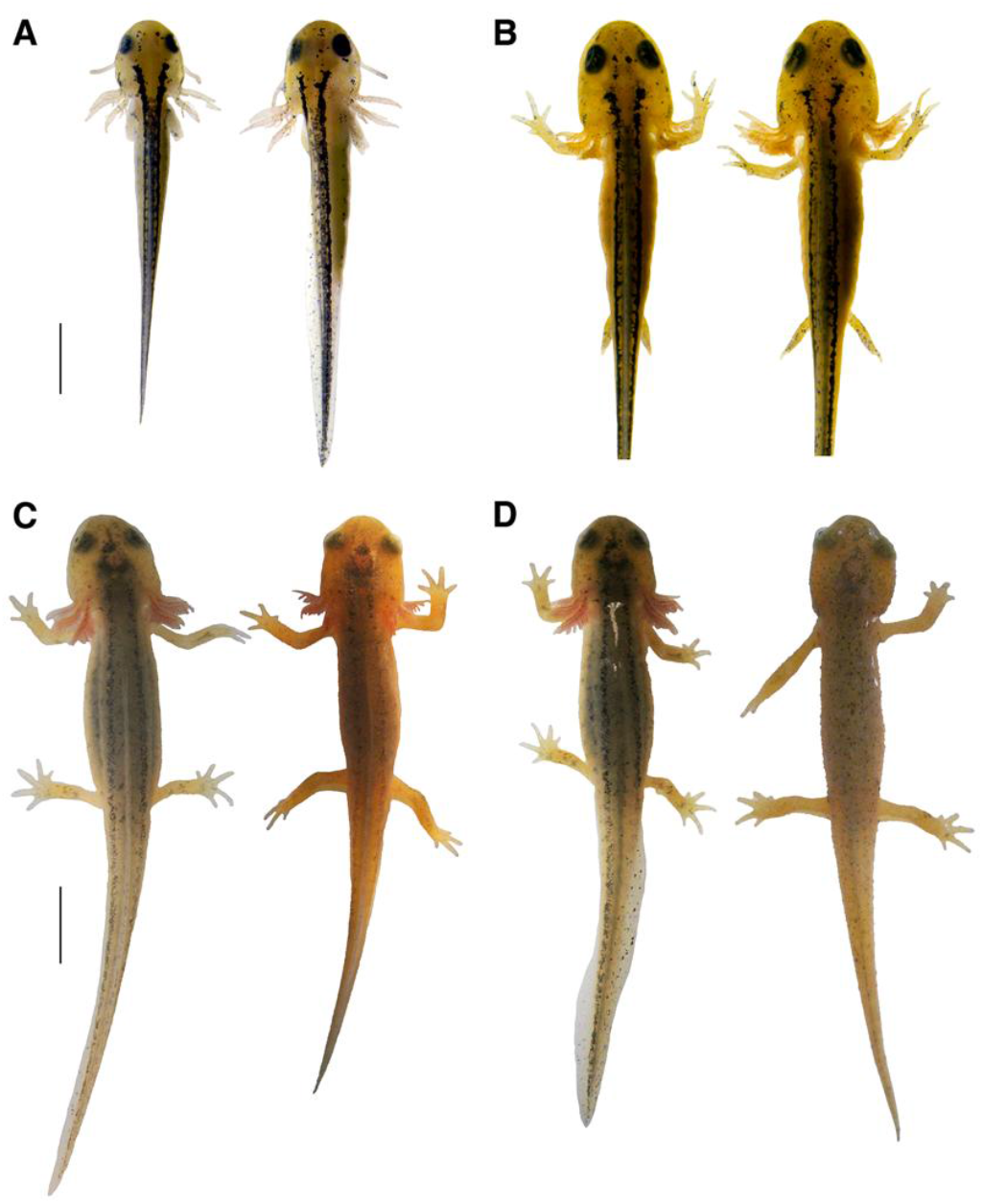
Differences in developmental rate in two animals from the same population, hatched on the same day and reared under the same conditions. In all panels, larva 5 is shown on the left and larva 4 is on the right. At hatching (**A**) larva 5 is at Stage 1 and larva 4 is at Stage 2. At 28 days post-hatching (**B**), both larvae are at Stage 8. At 105 days (**C**), larva 5 is at early Stage 12 and larva 4 is at mid-Stage 12. At 109 days (**D**) larva 5 is still at early-Stage 12 and larva 4 has progressed to Stage 13. The scale bar for the top row = 2 mm and that for the bottom row = 5 mm.

By the end of the study, the least developed subject, larva 5, had reached mid-Stage 12. The larva lingered in early Stage 12 for 45 days while growing in size. The larva entered mid-Stage 12 five days before it died, at which point it still had fluffy gills, visible yet receding skin stripes, and only mildly rough skin. The other larva had progressed to late Stage 12 and Stage 13 several days earlier, possessing rough skin and post-metamorphic pigmentation.

## Discussion

Newts of the genus *Taricha* have a rich history of use as models for studying both amphibian embryonic development and chemical defenses [13]. Early studies launched years of investigation into *Taricha* toxicity via a potent neurotoxin, tetrodotoxin (TTX, [14]). The discovery of TTX-producing bacterial symbionts living among the skin microbiota of these newts [15] generates further questions about the acquisition and maintenance of TTX in these animals. Dramatic changes associated with early development and metamorphosis, including changes in skin composition [16], hormonal cues, and immune system development [17] influence the formation of the early microbiome. Such early transformational cues may therefore also shape future toxicity of adult individuals. A standardized approach to staging will help to integrate information across studies for a more comprehensive understanding of the dynamics that shape the life history and toxicity of newts.

We identify 13 discrete larval stages, using limb development and balancer and gill recession as reference points while also describing more gradual feature changes. Through our observations, we were able to link particular developmental stages and anatomical changes. Hatchling larvae possess a visible yolk sac and they do not eat for several days. In parallel with the reduction of the yolk sac, the larvae transition to feeding on live prey. The balancers serve as temporary tools for navigating their environment, receding at Stage 5, when their forelimbs are developed enough to facilitate locomotion and stability. During mid-Stage 12, as the gills begin to reduce in size, the juveniles spend an increasing proportion of their time out of the water, suggesting that in the wild these animals may be becoming terrestrial and initiating their first migration out of their aquatic habitats at this stage.

With our staging table, we can also correlate past histological studies with the macroscopic changes in texture and pigmentation that shape developing larval skin. At hatching (Stage 1-2), the skin is smooth, sparsely pigmented, and translucent in places, particularly along the ventral side of the animal. The migration of pigment cells to form dorsal stripes is described by Tucker and Erickson [18] as beginning during embryonic development. Pigment cell migration is directed by the extracellular matrix and results in xanthophores occupying a large portion of the dorsal fin in early larval stages, while melanophores are sparsely present in the dorsal fin until the hindlimbs are partially developed (Stage 7). As the animal progresses through Stage 12, the skin becomes more opaque, likely corresponding to a restructuring of the histological composition of the skin [16]. Hatchlings have thin epidermal layers and minimal pigmentation, rendering their internal anatomy visible. The epidermis becomes thicker and more densely populated with melanocytes and keratin as the larvae approach metamorphosis. As larvae enter metamorphosis during Stage 12, the skin surface also develops distinct bumps associated with pores leading to newly developed dermal glands, including the granular glands in which *Taricha* store TTX and antimicrobial peptides [19].

Patterns of salamander development can vary from species to species and will not be captured by a singular representative development table. Species pigmentation development patterns or possession of balancers are correlated with whether the larvae hatch in ponds or streams, and, in the case of balancers, the extent of forelimb development at hatching [20, 21]. As summarized by Cao *et al*. [22], patterns of digit development can also differ across species. The initial digit primordia may arise in pyramidal, bifurcated, or triforked patterns at the distal end of the forelimb. The degree of limb development upon hatching can also differ greatly among species. While *Taricha torosa* hatches with either no digits or at most digit primordia present as a bifurcation of the distal forelimbs, larval *Hemidactylium scutatum* hatch with four digits present on the forelimb and mobile forelimb joints. Along with these well-developed appendages, *H. scutatum* larvae lack balancers throughout their development [23].

Further, even under controlled conditions, rates of development within a species can be highly variable. In agreement with the results from Hopkins et al. [24], our larvae hatched at varied stages of the Harrison development table, at which point digit primordia may or may not be present on the forelimbs. Maternal effects and predator cues can both influence level of development upon hatching as well as larval developmental time before metamorphosis [25, 26]. Each larva in our study developed at a different rate, demonstrating that even with consistent temperature, water quality, and food availability, time from hatching is not a dependable method for standardizing results.

Finally, while a staging system based on discrete feature changes is beneficial, we also describe more incremental developmental changes that may span multiple stages. Many of these more gradual changes occur during the longest stage, Stage 12, where the animal enters metamorphosis. These incremental morphological changes can indicate the onset of important molecular processes: for example, changes to the skin texture indicate major restructuring of the skin and may indicate the onset of skin gland development. Therefore, noting the texture of the skin in addition to labeling a larva as a Stage 12 individual can provide greater context. Continued creation of diverse post-embryonic staging tables such as the one described here will contribute to a strong foundation for integrating information from diverse studies on amphibian ecology, toxicology, microbiology, and more.

